# Spatially-Aware Clustering of Ion Images in Mass Spectrometry Imaging Data Using Deep Learning

**DOI:** 10.1101/2020.09.25.285619

**Authors:** Wanqiu Zhang, Marc Claesen, Thomas Moerman, M. Reid Groseclose, Etienne Waelkens, Bart De Moor, Nico Verbeeck

## Abstract

Computational analysis is crucial to capitalize on the wealth of spatio-molecular information generated by mass spectrometry imaging (MSI) experiments. Currently, the spatial information available in MSI data is often under-utilized, due to the challenges of in-depth spatial pattern extraction.

The advent of deep learning has greatly facilitated such complex spatial analysis. In this work, we use a pre-trained neural network to extract high-level features from ion images in MSI data, and test whether this improves downstream data analysis. The resulting neural network interpretation of ion images, coined *neural ion images*, are used to cluster ion images based on spatial expressions.

We evaluate the impact of neural ion images on two ion image clustering pipelines, namely DBSCAN clustering, combined with UMAP-based dimensionality reduction, and k-means clustering. In both pipelines, we compare regular and neural ion images from two different MSI datasets. All tested pipelines could extract underlying spatial patterns, but the neural network-based pipelines provided better assignment of ion images, with more fine-grained clusters, and greater consistency in the spatial structures assigned to individual clusters.

Additionally, we introduce the Relative Isotope Ratio metric to quantitatively evaluate clustering quality. The resulting scores show that isotopical m/z values are more often clustered together in the neural network-based pipeline, indicating improved clustering outcomes.

The usefulness of neural ion images extends beyond clustering towards a generic framework to incorporate spatial information into any MSI-focused machine learning pipeline, both supervised and unsupervised.

## Introduction

Mass Spectrometry Imaging (MSI) is a powerful, label-free molecular imaging technology that enables mapping the spatial distribution of thousands of biomolecules in a tissue section, based on a single experiment [13]. Due to its ability to combine rich biochemical characterisation with spatial information, MSI technology is rapidly being adopted for a panoply of applications, including biomarker discovery, clinical diagnostics and drug delivery studies [17,34,35]. While a number of different MSI variants exist [11], in general MSI operates by first overlaying the tissue with a virtual rectangular grid, and then collecting a mass spectrum at each grid location. Each of these collected mass spectra is a histogram of biomolecular ions counts, partitioned by their mass-to-charge values (*m/z*), within a target mass-to-charge range. MSI experiments result in a three dimensional data cube, with spatial coordinates (*x* and *y*) and a *m/z* axis containing the spectral information. Ion images are constructed by plotting the intensities for a single mass bin (*m/z* value) for each acquired pixel, i.e., (*x, y*) grid location in the tissue.

Technological advancements continue to push the capabilities of MSI, leading to improvements in specificity, sensitivity and speed, which have translated into the detection of larger numbers of biomolecular species at ever-increasing spatial resolution. A corollary of these technological improvements is that MSI datasets have grown substantially over the years. At the time of writing, a single experiment commonly generates tens of gigabytes of raw data, but this can even range into terabytes. Moreover, MSI data analysis is encumbered by the large number of variables measured, in terms of pixels, *m/z* bins or both.

Unless prior targets of interest are known, it is infeasible for a researcher to manually investigate and find molecular differences in these large datasets, and, as a result, computational approaches have become indispensable to support data analysis. A wide variety of unsupervised and supervised machine learning methods have been used in a broad range of applications. In this paper, we focus on unsupervised learning methods, which support exploring the underlying patterns within the MSI data, providing open-ended exploratory insights in the data. These methods include factorization methods (i.e. principal component analysis, nonnegative matrix factorization) and clustering methods, e.g. to group spectra or pixels with similar chemical expression into the same cluster [40]. Supervised learning methods, on the other hand, are used when prior knowledge or annotations are available, such as target *m/z*’s, or regions of interest, which can be used to guide the analysis, and, for example, extract differentially expressed molecular ions between regions in the tissue.

Using appropriate *distance measures* can substantially improve the outcomes of both supervised and unsupervised data analysis. A distance measure formalizes how to quantify similarity between data instances, e.g., between spectra originating from different pixels or between ion images associated to different *m/z* values. A recurring task in MSI data analysis is identifying which ions are co-localized, which translates into the problem of clustering *similar* ion images. As mentioned previously, such tasks are ideally tackled in a largely automated fashion to deal with the large amount of information in each MSI experiment. In order to answer such questions, a human expert would implicitly rely on his or her excellent visual pattern recognition abilities, and will use morphology, anatomical structure and saliency to assess the ions’ expression.

Unfortunately, it is difficult to translate the extensive spatial pattern recognition that is so easily performed by humans into robust, automated data analysis pipelines. To this day, most data analysis pipelines in MSI forgo the inclusion of available spatial information altogether [40]. Pipelines that do account for spatial information typically use measures such as spatial correlation or cosine distance to quantify similarity between ion images. As these are global measures, they are prone to miss relevant, localized differences between ion images. MSI machine learning methods that have explicitly focused on including spatial information into the analysis, generally focus only on either low-level or high-level information (e.g. through the inclusion of local pixel neighborhood information [2, 7] or the inclusion of anatomical information through anatomical atlases [41], respectively).

In this work, we propose a simple yet powerful approach to replicating human visual pattern matching. Owing to its simplicity, our approach can be incorporated in a wide range of data analysis pipelines, enabling them to exploit spatial information with few to no modifications. Our approach is rooted in deep learning, which is a class of models and associated learning methods based on artificial neural networks (ANNs). ANNs consist of layers of neurons, which are simple mathematical functions capable of doing basic, non-linear transformations of their inputs. By composing and stacking many layers, a *deep* network becomes able to recognize complex patterns in data, despite consisting of simple building blocks. This layered information processing resembles the workings of the brain, starting from low-level signals in initial layers up to high-level, domain-specific feature recognition in deeper layers.

In the case of computer vision tasks, the initial layers of the network act as image feature extractors, whereas deeper layers make higher level abstractions over those detected features [38]. In recent years, deep learning has shown tremendous potential in visual recognition tasks, having outperformed the state of the art machine learning algorithms in many computer vision tasks, and even human experts in certain biomedical tasks [19,33]. The strength of deep learning methods is their ability to learn and combine both low-level and high-level features and abstractions from image data. Deep learning has also previously been used in the context of MSI without the inclusion of spatial information [5, 20, 39].

While deep neural networks are very powerful, the disadvantage is that they are complex, and have a high number of parameters to be learned, and as such require a huge amount of - ideally labeled - data to learn powerful, high level abstractions. Directly learning those parameters from scratch every time for new datasets poses a big computational challenge, and is prone to overfitting in a setting such as MSI, where a lot of data is available on individual tissues, but the number of different tissues measured in a study is often limited, which is required to achieve good generalization. While it is possible to mitigate the issue of overfitting to a certain extent using data augmentation, for example by applying transformations to the original data [24] or by using generative adversarial networks [3,16,31] to automatically generate new representative data.

Moreover, deep learning is highly conducive to transfer learning, in which models trained for a certain task are re-purposed for other, related tasks [8]. In transfer learning, the requirement that the training data must have the same distribution as the test data is relaxed, i.e., a model is trained on one task, and the learned model is (fully or partially) transferred to another task. The more similar the task is, the better this approach generally works. Model-based transfer learning is particularly powerful for computer vision applications [23], because a lot of implicit knowledge, i.e., general image feature recognition, is shared amongst most practical tasks. Oquab et al. [28] have shown that the front layers of a convolutional neural network, that has been pre-trained on a large-scale annotated dataset, can be efficiently transferred to another computer vision task where limited data is available. Furthermore, the researchers noted that the way the neural networks interpreted the images was reminiscent of a human operator. A number of other studies have shown similar potential for re-purposing of neural networks for various downstream tasks [21,25].

In this work we will use the well-known Xception network [14], which has been trained on the ImageNet database [24] consisting of over a million images. Due to the large variety of the images included in this dataset, networks that perform well on this dataset, perform well on many other computer vision tasks [23], and as such can be used as general purpose, image feature extractors. Xception, for example, has previously been successfully used for feature extraction in various life science applications such as microscopy and electron microscopy 32, 42]. In the context of MSI, we will pass each ion image in the dataset through the network Xception, which generates a vector representation of each image, that encodes the high-level features within the image.

Mathematically, we use the neural network as an embedding function for ion images. The aim of this strategy is to map images with a high visual similarity close together, largely in line with how a human observer would interpret spatial resemblance between the original images. We will call these vector representations *neural ion images*, to highlight their origin and clearly contrast them with regular ion images. Note that these neural ion images are abstract, high-dimensional representations without any direct visual interpretation.

Ovchinnikova et al. [29] have recently also used Xception for evaluation of MSI ion image co-localization. Here, the Xception model was used to compare the visual ranking of ion image similarity by human experts to those produced by various distance measures and algorithms, such as the Xception network. However, here the network was used in a supervised way, i.e it was further trained using labels provided by human experts. The algorithm showed good performance in this task, albeit similar in performance to standard cosine distance for their use case (assessing the algorithm over different datasets). In contrast, our approach uses the pre-trained Xception model in a completely unsupervised way, without any domain transfer or fine tuning, and as such we show that it is directly usable for a wide breadth of applications in MSI, regardless of how much domain-specific data or computational resources are available.

It is important to stress that the resulting neural ion images can conceptually replace regular ion images as inputs for most downstream machine learning and statistical analyses, both supervised and unsupervised. Using neural ion images enables improving existing data analysis pipelines to better incorporate spatial expression patterns with minimal change to the full pipeline.

In this work, we focus on assessing the merits of neural ion images for the unsupervised task of clustering, i.e. grouping together, ion images based on their spatial expression. Similar to how factorization is often used in the context of MSI, the clustering of ion images reveals the different spatial patterns that are present in a MSI dataset. This is done by visualizing the mean ion image for each of the different clusters (or groups) of ion images that are found in the dataset. Furthermore, this method provides direct insight into which *m/z* values exhibit comparable spatial expressions, compared to factorization where this relationship is not always straightforward [40]. Alexandrov et al. [1] previously applied a probabilistic clustering algorithm (a Gaussian mixture model) to cluster ion images in MALDI MSI datasets based on their spatial similarity. Similarly, by Konicek et al. [22] used k-means to cluster together ion images in a TOF-SIMS dataset.

Here, our goal is to improve on this work by integrating the pre-trained Xception model into the clustering pipeline, and as such find clusters with a higher specificity, based on localized spatial features, and with a greater consistency in the images assigned to the same cluster.

## Methods

### Materials

We demonstrate our proposed method on two MSI datasets, namely one from human lymph node and one from a mouse kidney tissue.

The first dataset is collected from a human lymph node tissue sample using the Bruker rapifleX MALDI Tissuetyper. The tissue sample was removed and snap frozen in liquid nitrogen (−80 *°C*). Tissue sections of 7 *μm* thickness were acquired using a cryostat and thaw-mounted onto ITO-coated glass slides, after which 2,5-DHB matrix was deposited by sublimation. Experiments focused on the 620 to 1200 *m/z* range, using a sampling resolution of 10 *μm*, collecting roughly 500.000 pixels and 8.000 ion images for this tissue.

The second dataset is collected from a mouse kidney tissue section. The tissue sample was snap frozen in liquid nitrogen at necropsy. Tissue sections were collected in a cryostat (CM3050S, Leica, Buffalo Grove, IL; chamber temp. −20°*C* and object temp. −18°*C*) at 10 *μm* thickness and thaw-mounted onto ITO-coated glass slides. 2,5-DHB matrix prepared at 25 mg/mL in methanol:water (1:1, v:v)(0.5% TFA) was applied to the tissue sections using a TM Sprayer (HTX Technologies, Chapel Hill, NC) automated spray device. The following parameters were used to achieve a target matrix density of 0.4938 *mg/cm*^2^: flow rate 0.05 *ml/min*; *N*_2_ Pressure 10 *psi*; Spray temperature 70 °*C*; velocity 1350 *mm/min*; track spacing 3 *mm*; track offset 1.5 *mm*; 16 passes. MALDI MSI was conducted at a sampling resolution of 50 *μm* on a Bruker scimaX MRMS. Fullscan data was acquired from *m/z* 200-1200 using 250 laser shots per pixel at a frequency of 2 *kHz* and an estimated resolving power of 66K at *m/z* 400. This study was conducted in accordance with the GSK Policy on the Care, Welfare and Treatment of Laboratory Animals and was reviewed by the Institutional Animal Care and Use Committee either at GSK or by the ethical review process at the institution where the work was performed.

### Ion Image Clustering

In order to assess the merits of our deep learning approach, and to demonstrate how it can be easily integrated into any data analysis pipeline, we will investigate two different, well-known clustering methods, both with and without the use of neural ion images. Figure 1 gives an overview of the ion image clustering workflow, and shows three separate processing blocks, which will be discussed in greater detail below. In the first block, ‘‘Shared preprocessing”, all images of the MSI dataset receive a common preprocessing treatment. The second block, “Generation of neural ion images”, contains the neural network steps, and either passes the ion images through the pre-trained neural network to generate “neural ion images”, or leaves the images unaltered (“regular ion images”). In the third and final block, “Clustering steps”, the clustering of the images is done using k-means or a combination of UMAP and DBSCAN (U-D for brevity), and takes as an input either neural or regular ion images. This results in a total of four different ion image clustering pipelines, which we will refer to as regular U-D, neural U-D, regular k-means, and neural k-means.

**Figure 1.**
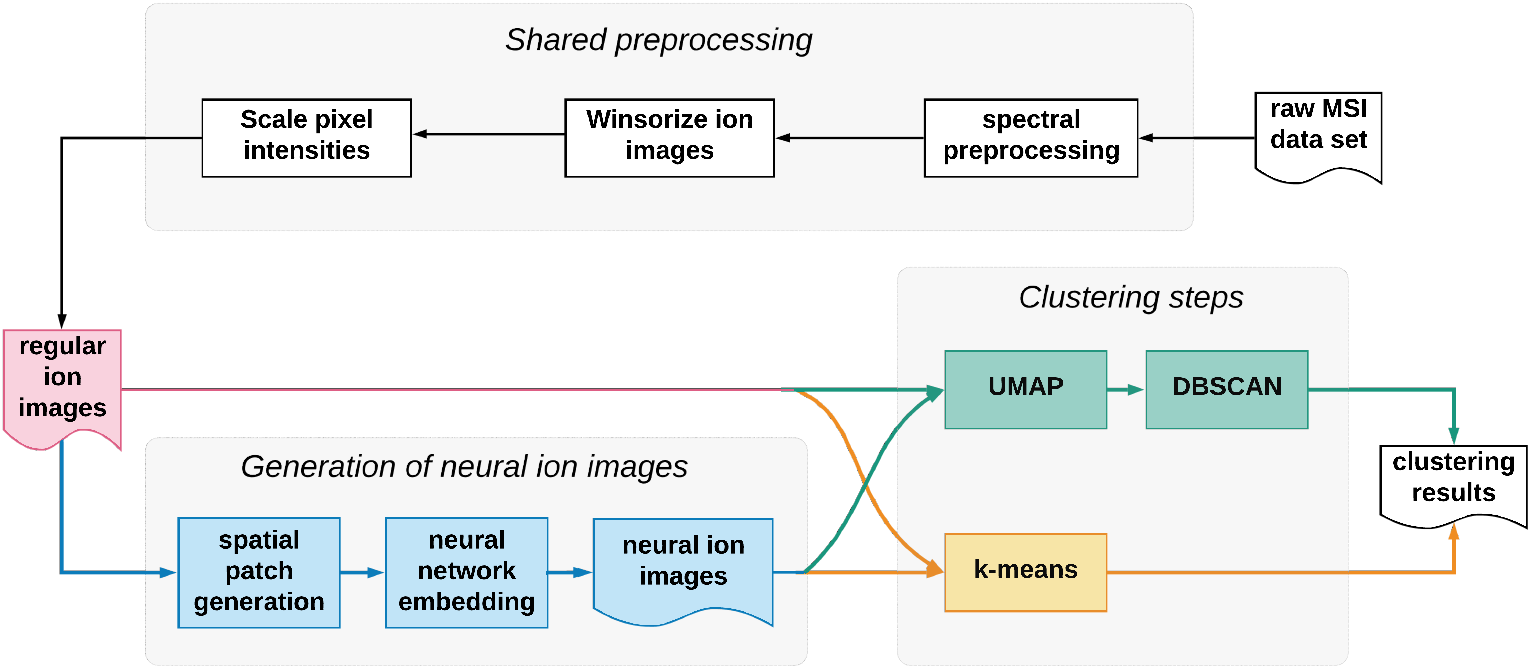
Pipelines: the top panel contains a series shared preprocessing steps. The lower left panel shows the additional steps (in blue) from our proposed method compared with regular clustering pipelines (in red). The additional steps from our proposed model include spatial patch generation, neural network embedding and neural ion images access. Finally two different clustering methods are conducted, which are DBSCAN (after UMAP dimentionality reduction) and k-means.

### Shared preprocessing

Before applying any clustering pipeline, we perform basic preprocessing on the raw spectra and, afterwards, outlier removal and standardization on the resulting ion images. These procedures are done separately for each dataset.

1. *Spectral preprocessing* The lymph node data is normalized based on Total Ion Current (TIC) and baseline corrected using median filtering. The mouse kidney data is normalized based on TIC and peak-picked. Peak picking was conducted using a S/N threshold of 5 and the Gibbs, harmonic, and magnetron peaks were filtered during acquisition. The SQLite peak list files were processed in MATLAB using custom software to recalibrate each spectrum using a linear best fit model and then bin corresponding peaks across all spectra in the dataset to a common *m/z*. The binned data was then converted to imzML for further processing.
2. *Ion image standardization* The ion images for both datasets are individually winsorized and then standardized to a common range prior to clustering.

Winsorizing is a well known robust statistical estimation technique which limits extreme values to reduce the effect of potential outliers [18,43]. Winsorizing involves clipping extreme values, usually symmetrically on both extremities of a distribution, before computing location statistic(s) of interest such as the mean. In our case, we only winsorized above the 95th percentile to reduce the impact of high intensity outliers commonly found in MSI data. Winsorizing the lower extremity is not necessary for spectral data since there is an explicit limit at 0 intensity. After winsorizing, the resulting pixel intensities for each image are scaled to the range [0,1]. The resulting preprocessed images will henceforth be referred to as *regular ion images*, and serve as input for the next stage of the pipeline.

### Generation of neural ion images

Our proposed approach involves a couple of extra steps to convert regular ion images into neural ion images. Generating neural images from individual regular ion images is done as follows:

1. *Patch generation* A neural network typically requires images of certain shape as inputs. Since ion images can be of any shape, we first define a set of overlapping patches across the ion images, which decouples our pipeline from any specific shape of ion images.
2. *Generating patch embeddings* The pixel data for each patch is fed into the neural network as a single data instance, which results into an associated neural embedding per patch.
3. *Aggregating patch-level embeddings* The neural ion image is constructed by aggregating the embeddings of individual patches into a single vector representation for each regular ion image.

We discuss each of the aforementioned steps in more detail below.

#### 1. Patch Generation

Ion images tend to be larger than the default input size expected by neural networks. For example, the Xception network we used [14], accepts square patches of 299 × 299 pixels by default, with a minimum of 71 × 71 pixels. These shape mismatches can be fixed via patch generation, which is especially common when using convolutional neural networks to analyze high-resolution images. Patch generation means that, rather than providing the full image to the network, the image is instead split up into smaller “patches” that are then given as an input to the network, thus preserving all the details in the original image in one or several patches. An example of patch generation is shown in Figure 2. The size of the patches is important as it determines the context that the network “sees” in one go. Therefore, we tested different patch sizes, as described in the Results section.

**Figure 2.**
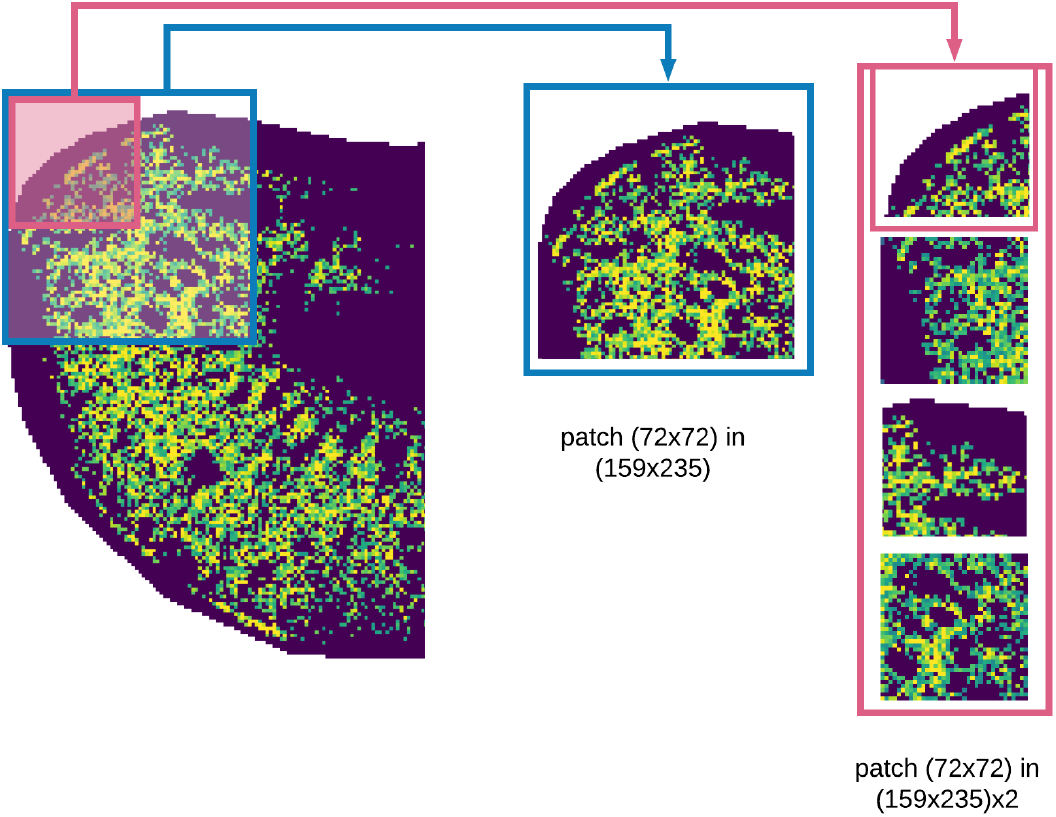
From left to right: a regular ion image from the mouse kidney dataset, generated patches with different sizes. In order to get the ideal size of patches, we up-sample the full ion image before splitting into multiple patches. All results were obtained using up-sampled patches, as indicated in pink, which have a physical size of roughly 1.85 × 1.85 mm.

#### 2. Generating patch embeddings

A neural network is used to generate a vector representation (embedding) of each patch, which ideally captures all relevant morphological information present in the patch. The network itself is pluggable in our pipeline, so other networks could also be used. We leveraged the widely used Xception model because it has shown strong performance on recent benchmarks [14], despite being relatively small compared to other models with only 36 convolutional layers. Owing to its small size, its inference speed is higher then that of many other models with comparable accuracy [10]. Finally, by using a pre-trained network, we avoid the demanding task of designing and training the network, which would require a lot of data as well as computational resources. We use the Xception network as is available in the Keras Python package https://keras.io/.

#### 3. Aggregating patch-level embeddings

Finally, after obtaining an embedding associated to each patch, these must be integrated into a final vector representation that constitutes the neural ion image. We opted to use max-pooling because the neural ion image then essentially captures whether a spatial pattern (single output feature in all patch embeddings) was detected in *any* of the patches. This optimizes the neural ion images’ sensitivity to detect localized features.

### Clustering steps

We use two different clustering approaches in this work, namely DBSCAN, combined with UMAP, and k-means clustering. As both of these clustering pipelines expect vector inputs, regular ion images were reshaped to vectors (neural ion images are already vectors). In this work, we purposefully selected two fundamentally different clustering methods to properly assess the merits of using neural ion images instead of regular ion images.

#### UMAP-DBSCAN

The first approach, DBSCAN, which stands for Density-Based Spatial Clustering of Applications with Noise [15], is a popular density-based clustering algorithm that inspired several extensions (i.e. HDBSCAN [12,37]). DBSCAN aims to find high density regions in feature space where samples are closely packed, i.e. samples that have a lot of other closely resembling samples (neighbors). Based on a user-defined radius, the algorithm estimates a minimum density level (i.e. minimum number of neighbors within a radius). Clusters are then regions where groups of samples exceed this density threshold, whereas points that are in low density regions are put in a common *noise cluster*. DBSCAN is widely available in different implementations, here we use the scikit-learn implementation [30].

Given that we are clustering ion images, each pixel can be seen as a separate feature, or measured variable, and, as such, the number of features per image is very high (e.g. ~ 500.000 pixels per regular ion image for the lymph node dataset, thousands of variables per neural ion image). High-dimensional feature spaces are inherently sparse due to the curse of dimensionality [6], which encumbers the process of identifying clusters based on density [40]. It is therefore advisable to first do a dimensionality reduction step prior to performing DBSCAN, which we did using Uniform Manifold Approximation and Projection (UMAP) [26]. Briefly, UMAP aims to form a topological representation of the original high-dimensional data by finding local manifold approximates using their local fuzzy simplicial set representation. Prior work has shown promising results of applying UMAP on spectra in MSI data [36] (as opposed to ion images as we are doing here).

In this paper we used UMAP to reduce the dimensionality of regular and neural ion images to three prior to applying DBSCAN. Mapping to three dimensions enables visualizing the resulting embedding, which helps in interpreting the results, and is sufficiently low-dimensional to effectively support DBSCAN. Different parameters were manually optimized for UMAP and DBSCAN for the different models because the inputs for each pipeline are very different. To facilitate a fair comparison, we did a range of experiments and selected the most promising model with selected parameters for each approach. For the regular ion images, each ion image was reshaped to a vector of size one-by-’number of pixels’, and cosine was used as the distance measure between the resulting image vectors in the UMAP algorithm. For the neural ion images, similarly the cosine distance was used as the distance measure between the neural ion image vectors. The Python code by UMAP’s creator was used [27]. For brevity, we will refer to combination of UMAP-DBSCAN as U-D.

#### k-means

The second clustering algorithm used in this paper is the well-known k-means clustering [9], which is available through scikit-learn [30]. k-means is a highly popular clustering algorithm used in a wide variety of applications owing to its simplicity and speed [4]. Conceptually, it tries to separate objects in a number of groups with low intra-group variance by minimizing the squared distance between samples and the prototype vector (mean) of the cluster they are associated to. The Euclidean distance (standard in k-means) was used for all experiments.

#### Practical differences between U-D and k-means

In contrast to DBSCAN, k-means requires the number of clusters to be decided in advance, as we will discuss further below. The ability to explicitly reject certain samples from clustering grants DBSCAN increased robustness to noise and outliers compared to k-means, which always forces each sample into a cluster. Conceptually, owing to its nonlinear nature, U-D is expected to identify more subtle trends within data, but requires more expertise to optimize its various tuning parameters. From a computational perspective, k-means is significantly faster.

## Evaluation of Clustering Results

We proposed different methods to evaluate the quality of the resulting clusters from the different pipelines.

### Number of Clusters

After clustering, we first check the number of meaningful clusters. Meaningful clusters should not include the noise cluster as defined by DBSCAN (as previously discussed) or clusters that contain a single image when using k-means. Furthermore, we also consider clusters that only contain noisy images to be non-meaningful clusters. This is especially relevant in the lymph node data, where no peak-picking was done, and as a result a lot of noisy images are expected in the dataset. We assume that, the more meaningful clusters are extracted, with distinct mean images, the more likely it is that more underlying structures are detected from the dataset.

### Relative isotope ratio (RIR)

In order to objectively compare the performance of different clustering pipelines, we propose a metric that quantifies how well a clustering groups ion images stemming from isotopic *m/z* values. Two *m/z* values are considered isotopes if and only if the following criteria are met:

1. The spectral distance equals 1.003m/z±*δ*, where *δ* ≠ 0 allows for small deviations in mass-to-charge values. The value of *δ* depends on the mass accuracy of the MSI data. This spectral distance is primarily intended to identify carbon isotopes, however if *δ* is set large enough this will capture other isotopes as well.
2. The corresponding ion images should exhibit a clearly similar spatial expression, which we assess using Pearson correlation. Specifically, the correlation between both ion images should be at least 0.85.

To assess the quality of isotope grouping for a given clustering, we count the number of isotopes that are correctly grouped in the same clusters and divide this count by the total count of isotopes in the dataset. This fraction *F* ∈ [0,1] captures how many of the isotopes in the dataset are clustered together (higher is better). However, this fraction favors large clusters, because then the probability of clustering isotopes together increases. This can intuitively be seen by the trivial edge case of having a single cluster, in which all isotopes are obviously clustered together, thus yielding a seemingly perfect score.

As such, because the *expected* value of *F* depends on the size of the clusters in a clustering, we benchmark the isotope fraction of a given clustering (*F_clust_*) to that of a simulated random clustering with the same cluster sizes (*F_random_*). This random clustering is simulated in a bootstrap fashion to get a consistent estimate of the expected isotope fraction. Our *relative isotope ratio* metric 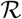 is the relative increase in isotope grouping for a given clustering compared to its random baseline, i.e., 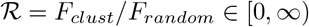, where higher is better. Any clustering that is better than random should yield 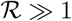.

### Visualization of Clustering Results

In order to visualize the spatial results, we compute the mean ion image for each cluster by first winsorizing each ion image from all clusters, followed by stratified averaging over all *m/z* images per cluster. Those mean ion images then show the various spatial patterns detected by different clustering pipelines. We also generate a mean spectrum for each dataset with clusters assignment for each *m/z* value.

Furthermore, for U-D clustering, we visualize the 3D embeddings after UMAP dimensionality reduction, which results in a 3D scatter plot where each *m/z* image is a point with clusters assignment. This scatter plot illustrates how all *m/z* images are split and clustered in different groups, the distance between different clusters, and the tightness of each cluster.

Finally, we utilize histological information from concomitant microscopy images to check whether the clusters are biologically relevant. These microscopy images were obtained on serial sections to the MSI data. To assess biological relevance, we construct a false color image from the mean ion images of 3 different clusters, where each image uses a separate color channel (RGB), which is then overlaid onto the microscopic image. If the overlay shows a clear co-localization of cluster images and morphological patterns in the associated microscopic images, it can be assumed that the clusters are biologically relevant.

## Results and Discussion

We start by preprocessing the data, as per the “shared preprocessing” block described in Figure 1. Spectral preprocessing differs for both datasets: the lymph node dataset is TIC normalized and baseline corrected, whereas the mouse kidney dataset is TIC normalized and peak picked. Next, each individual ion image is winsorized, substituting pixels with outlying intensity values, after which it is scaled so that pixel intensities lie between 0 to 1. After the shared preprocessing, we start one of four different clustering pipelines, namely UMAP-DBSCAN on regular ion images (regular U-D), UMAP-DBSCAN on neural ion images (neural U-D), k-means on regular ion images (regular k-means), and k-means on neural ion images (neural k-means). Each of the four clustering pipelines is used to process the human lymph node and mouse kidney datasets separately.

### Generation of neural ion images

Once the MSI data is preprocessed, the regular ion images can be used to create neural ion images using the pre-trained neural network. First, we need to generate image patches with a size appropriate for the pre-trained Xception network. As explained previously, the size of the patches is important, as this determines the context that the network receives as an input. We therefore aimed to select a patch size with a physical size that is large enough to capture morphological patterns and anatomical structures, testing various patch sizes.

We found that patches with sides of a physical size of around 1-2 mm work well in practice. For the lymph node dataset, at a 10 *μm* sampling resolution, this results in a patch size of 150 × 150 pixels, in a total image size of 1,183 × 696 pixels. For the mouse kidney dataset, we tried to get patches with a similar size, however, given the 50 *μm* resolution, a patch with the minimum pixel size of 71 × 71 for the Xception network would still be too large (~ 3.5 mm). We therefore upsample the ion images for mouse kidney data with a factor of two, so that we obtain more, smaller patches with a more appropriate physical size. Figure 2 shows a patch generation example from mouse kidney dataset, with the original patch sizes highlighted in blue and the patches after upsampling indicated in pink. Furthermore, it is important to note that we use overlapping patches to avoid edge artifacts, e.g., missing anatomical structure because it was located at the edge of a patch. In our patch generation procedure, we ensured a 40% to 50% spatial overlap between patches.

Using this procedure, we generate image patches for each ion image in the MSI dataset. The image patches are batched per ion image and are sequentially fed to the pre-trained network, which analyzes each patch and extracts low- and high-level image features from them. This translates to a vector per patch, that scores the various features detected by the network. The results for the individual patches are then combined in a max-pooling layer at the end of the network, which registers whether a spatial pattern is detected in any of the patches, to increase sensitivity for localized features. The result is a single vector (of size 2048) per ion image, which captures the “interpretation” of each ion image by the neural network, i.e., the neural ion image.

With both the regular and neural ion images available, we can use these as input for the clustering algorithms, in order retrieve the different spatial ion expressions in the dataset.

### Human lymph node dataset

#### UMAP-DBSCAN clustering results

We apply each of the four clustering pipelines to the human lymph node dataset, starting with the U-D pipelines. First we perform UMAP dimensionality reduction on both the regular and neural images, which maps the images from the original 500.000 (regular) and 2048 dimensional (neural) space respectively to a three dimensional embedding space. The goal here is to map similar images close together in this new embedding space. The resulting embeddings are shown in the scatterplots in Figure 3, where each point represents a *m/z* image.

This dimensionality reduction serves a double purpose, firstmost it serves as an important preprocessing step to facilitate subsequent clustering with DBSCAN, and second, mapping to 3 dimensions allows us to visualize the embeddings and get some insight into what is happening. We then use DBSCAN to find regions of high density (i.e. groups of images that are highly similar in the embedding space) in the data, retrieving 22 and 21 clusters for the regular and neural pipelines respectively, as show in table 1. The different clusters are assigned different colors in the scatter plot, and we see clear clusters of similar *m/z* images in the embeddings.

**Figure 3.**
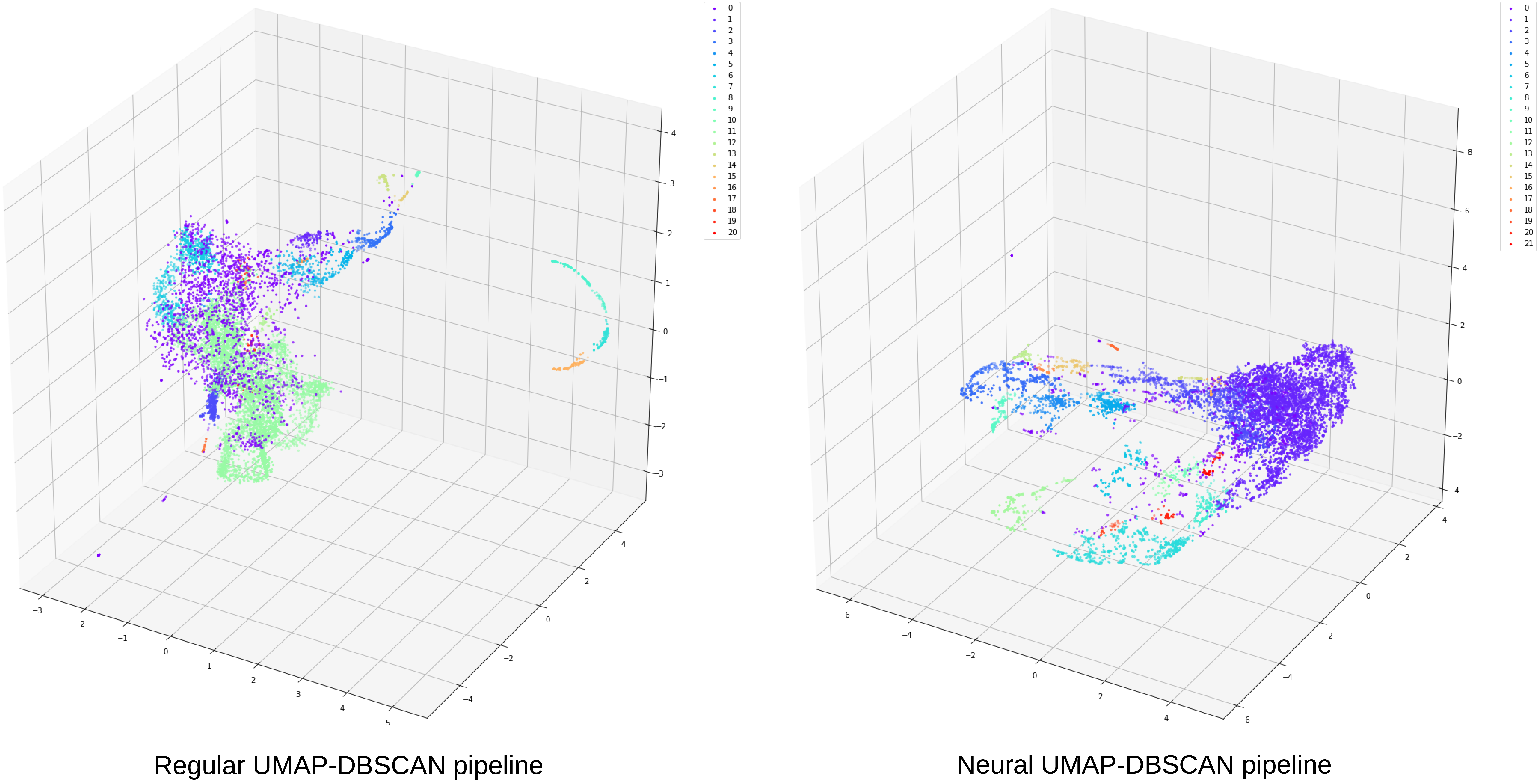
Scatter plots of resulting embeddings in 3 dimensional space after applying non-linear UMAP dimensionality reduction and DBSCAN clustering on the lymph node dataset. Each point represents a *m/z* image, with its colors showing cluster assignment as determined by DBSCAN. The left and right panels show clusterings on the regular and neural ion images respectively. The Neural U-D pipeline shows more localized and clearly defined clusters of *m/z* images than the regular U-D pipeline

**Table 1.** Overview of experimental results per configuration. RIR indicates relative isotope ratio (higher is better, best per dataset marked in bold). Noise images shows the size of the *noise cluster* identified by DBSCAN.

| Dataset | Clustering | Ion images | Clusters | RIR | Noise Images |
| --- | --- | --- | --- | --- | --- |
| lymph node | UMAP-DBSCAN | neural | 22 | **66.6** | 517 |
| lymph node | UMAP-DBSCAN | regular | 21 | 57.7 | 1671 |
| lymph node | k-means | neural | 20 | 64.5 | / |
| lymph node | k-means | regular | 20 | 56.2 | / |
| mouse kidney | UMAP-DBSCAN | neural | 40 | 35.4 | 188 |
| mouse kidney | UMAP-DBSCAN | regular | 21 | 20.4 | 63 |
| mouse kidney | k-means | neural | 60 | **35.6** | / |
| mouse kidney | k-means | regular | 20 | 23.4 | / |

Overall, the neural pipeline shows more localized clusters than the regular one. This is what we would expect, as the neural pipeline should make a better distinction of images based on localized extracted features by the neural network, than the more global cosine distance used in the regular pipeline. We also see a relatively large localized purple cluster in the neural U-D case, which is the noise cluster that DBSCAN defines (517 images); in the regular U-D pipeline this cluster is larger (1, 671 images, around 20% of total images) and much more spread out in the embedding space.

On inspection, the noise cluster in the regular pipeline shows a great number of images that still contain spatial structure, and were thus wrongly classified as noise. While the neural pipeline was not perfect, the noise cluster contains far less images with spatial structure. We note that the spatial resolution of this dataset is very high, but as a trade-off the mass resolution is relatively low. As such, even though the dataset is not peak-picked, many of the *m/z* bins contain spatial signal.

Overall, when comparing cluster assignments for the neural and regular pipelines, the neural pipeline showed a much better assignment of ion images, and a greater consistency in the spatial structure of images assigned to individual cluster, resulting in better, more representative mean cluster images. Figure 4 shows examples of the clustering of the lymph node dataset for the regular and neural clusterings. The bottom of the Figure shows the full mean spectrum with colors indicating assignments of *m/z* values to the clusters for neural and regular pipelines (top and bottom, respectively). A zoom-in for part of the spectrum is shown with detailed cluster assignments, showing marked ion images at the top. By their relative spectral distance of ~ 1 Da, and similar spatial distribution, we can assume that these marked images are probably isotopes, and as such we would expect all of these images to be allocated to the same cluster. In the left panel, in purple, we see assignments for each of these images for the regular pipeline. The images shown are the mean images of the cluster to which the different marked images are assigned. Firstly, we see that only two of the isotope images are assigned to the same cluster, whereas the other two images are assigned to different clusters, and secondly, we see that the mean images of the clusters that the images are assigned to, do not closely resemble the original isotope images. The neural pipeline, on the other hand, uses the features extracted by the neural network to correctly assign all images to the same cluster, with a mean image that closely resembles that of the original ion images, thus performing much better at this task. Furthermore, the mean cluster images of the neural pipeline show better definition of spatial structure, compared to those of the regular pipeline, which are generally much more blurry, indicating a more diverse collection of underlying images.

**Figure 4.**
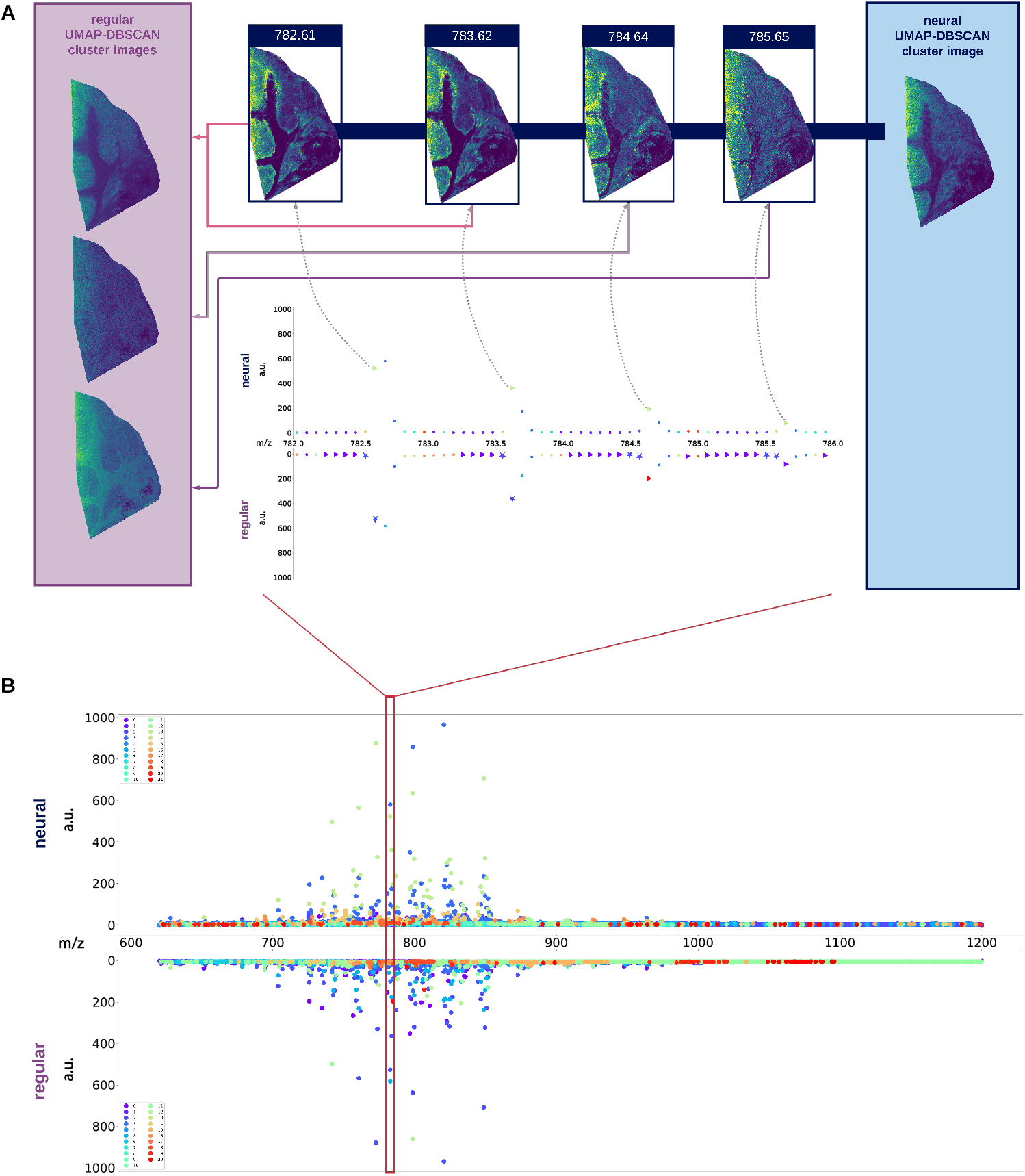
Ion image clustering results of the lymph node dataset. Panel B shows the full mean spectrum with cluster assignments for each *m/z* bin, for the neural (top) and regular (bottom) U-D pipelines. Panel A shows a zoom-in of the mean spectrum, highlighting an isotopic distribution, and four associated ion images. The purple (regular) and blue (neural) panels show the clusters (shown as mean images) to which the associated ion images are assigned. The neural pipeline shows better cluster assignments than the regular pipeline as (i) all isotope images are assigned to one and the same cluster, and (ii) its mean cluster image matches the associated isotopic ion images more closely than the regular pipeline.

This trend is observable in a large number of the clustered images, as is confirmed when we look at the Relative Isotope Ratio (RIR), in Table 1. This ratio measures the overall number of isotopes in a cluster compared to a randomized clustering, as explained in *Relative isotope ratio (RIR)*. The higher this number, the more isotope ion images are assigned to the same cluster, which is the expected behaviour. Looking at the table, we can see that the neural pipeline scores much better at this metric than the regular one, demonstrating the added benefit of adding a neural network-based interpretation layer to the pipeline.

Finally, we verify the clustering results by comparing the mean cluster ions with the stained H&E microscopy image of a neigboring tissue section. Figure 5 shows three example mean cluster images, originating from the neural U-D pipeline, on the right, highlighting different salient structures in the lymph node tissue. The center image shows a composite image overlaid on the microscopy, where each mean cluster image is assigned a different color channel, namely red, green and blue, that shows a clear overlap of the anatomical structure observed in microscopy with the biochemical information obtained from the MSI data. The green cluster image clearly shows the Germinal Centers (balloon shaped structures) found in lymphoid tissue. These are important areas for humoral immunity as, at these sites, activated B cells (B lymphocytes) accumulate and undergo further processing. The high-resolution, non-rigid registration was performed using Aspect Analytics’ proprietary registration pipeline.

**Figure 5.**
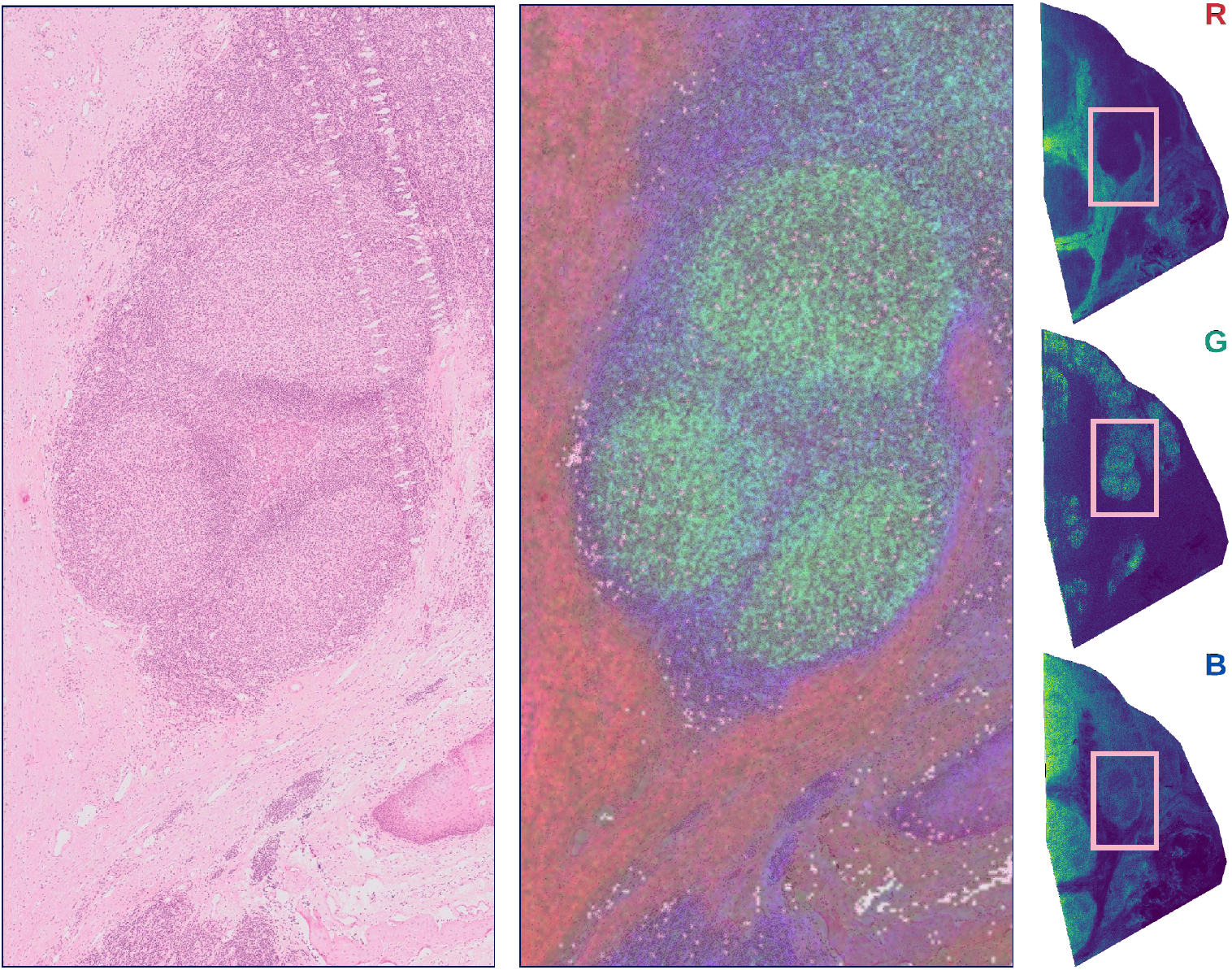
(Left) H&E stained microscopy image of neighboring lymph node tissue section. (Right) Three example mean cluster images for the neural U-D pipeline. (Center) Composite image combining the three mean cluster ion images overlaid onto the microscopy, showing clear co-localization of anatomical structures (via morphology) with the biochemical information obtained through clustering of the ion images from MSI.

#### k-means clustering results

Next, we apply k-means clustering directly on both the regular and neural ion images, as a baseline comparison, and an example of how neural images can be readily plugged in to existing algorithms. We experimented with different numbers of clusters, and found that, in line with the DBSCAN experiments, 20 clusters provided the best results in terms of detected meaningful clusters and RIR (see table 1). For the regular k-means pipeline, this results in 6 clusters which contain only a single image that had no clear spatial structure (and which we thus consider to be noise clusters). This is probably due to the high dimensionality of the data (500.000 pixels - features), and the sensitivity of k-means to noise. The neural pipeline did not show such single image clusters, even when increasing the number of clusters, probably due to the lower dimensionality of the neural images (2048) and a better grouping of the neural ion images due to the feature recognition by the neural network. When comparing the RIR, we see a similar trend to the U-D pipeline, namely an upregulation in the numbers of isotopes that are captured by the same cluster, when using the neural images over the regular ion images. This indicates that clustering results are better with the neural network input than without, which we also saw in manual inspection of the clusters.

### Mouse kidney data

In order to assess the general applicability of our methods, we also apply our clustering pipelines on a high mass resolution dataset obtained in mouse kidney, that has a significantly lower spatial resolution than the lymph node dataset, respectively 50 vs. 10 *μ*m.

As the spatial resolution affects the fine structure observed in the ion images, we want to see whether the neural network approach can improve clustering results similarly to the high spatial resolution lymph node dataset. As discussed above, we upsample the image data, so that patches are generated with a similar physical dimension to those in the lymph node data. We then pass the ion images through the neural network to perform feature extraction and generate the neural ion images, exactly as we did in the lymph node data.

### UMAP-DBSCAN clustering results

We first apply the U-D pipeline to the regular and neural ion images. As can be seen in Table 1, DBSCAN finds significantly more clusters in the neural (40) pipeline than the regular (21) one. Furthermore, similarly to the lymph node data, the RIR is significantly higher for the neural pipeline than the regular one, indicating that the retrieved clusters in the neural pipeline succeed in clustering together more isotopic ion images than the regular one, meaning that we not only get more clusters, but that these are also more relevant. When looking at the noise clusters, we see that this time the noise cluster in the neural pipeline is larger than the regular one, however the size is only 3.8% of the total number of ion images. We note that this time the data is peak-picked, and thus the noise cluster is much smaller.

The quality of the clustering is illustrated by the example shown in Figure 6, which shows three sets of ion images corresponding to three different isotopical distributions. Again, the blue panel shows mean cluster images for the neural pipeline, whereas the purple panel shows those for the regular pipeline. Similarly to the lymph node dataset, the neural pipeline correctly assigns images of the same isotope distribution to the same cluster. Contrary to the lymph node data, in this example, the regular pipeline also assigns the isotopic images to the same cluster. However, when comparing the mean cluster images to which the isotopic images are assigned, we see that, while not perfect, those of the neural pipeline match the isotopic images much closer than those of the regular pipeline. The neural pipeline succeeds in creating clusters with a higher specificity, which allows for distinguishing of the salient spatial distributions in the isotopic images at m/z 856.63 and 856.66. In contrast, the regular pipeline groups these patterns together, likely because it uses a more ‘‘global” distance measure that is prone to ignoring localized structure. More generally, we see a lot of these clusters that capture fine spatial structure, which are missing in the regular pipeline. Figure 7 shows how different mean ion images obtained with the neural U-D pipeline clearly differentiate distinct anatomical regions in the kidney, overlaying these on the H&E of a neighboring tissue section.

**Figure 6.**
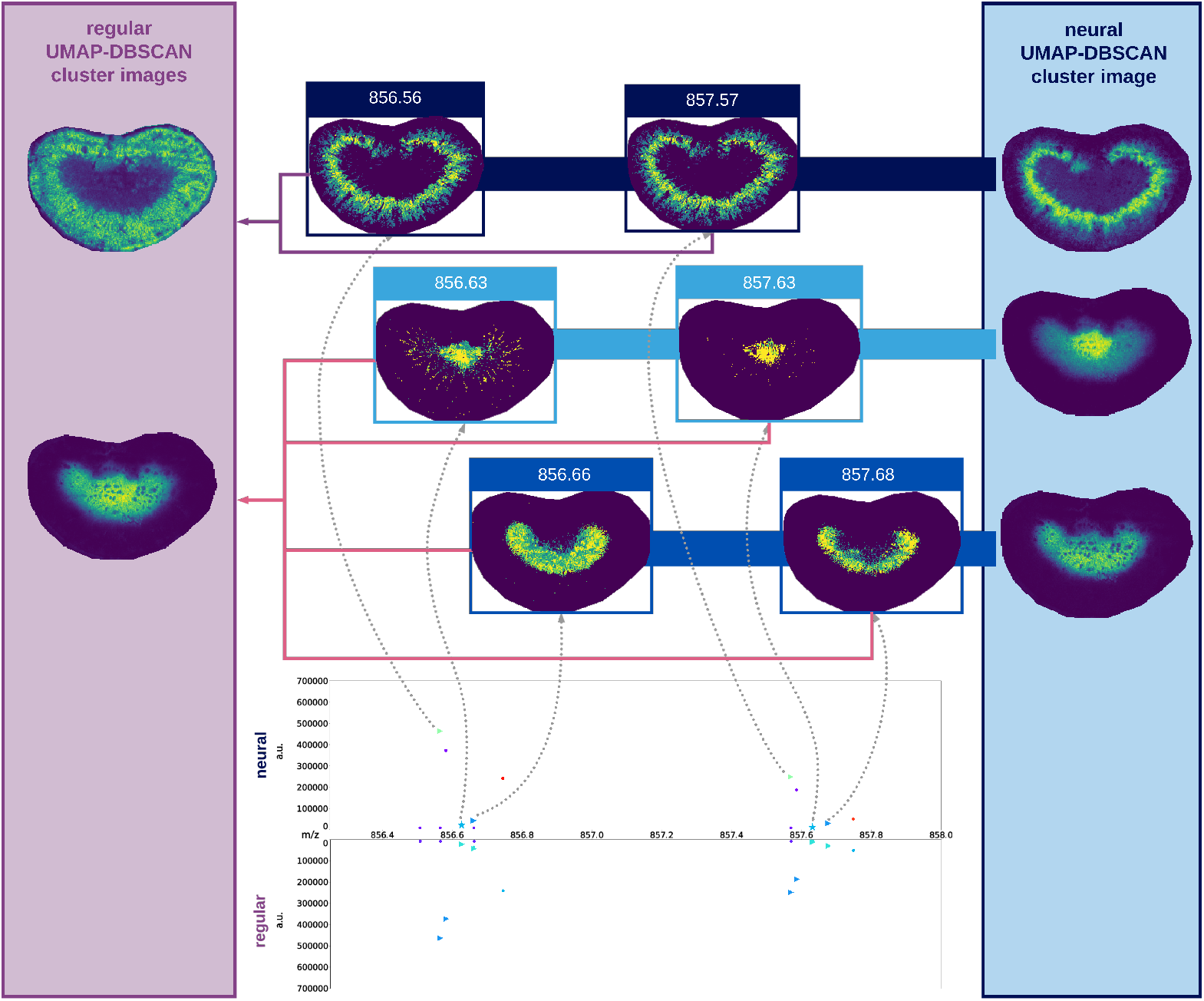
Ion image clustering results of the mouse kidney dataset. The Figure shows a zoom-in of the mean spectrum, highlighting an isotopic distribution, and marked associated ion images. Cluster assignments in the mean spectrum are shown for each *m/z* bin, for the neural (top) and regular (bottom) UMAP-DBSCAN pipelines. The purple (regular) and blue (neural) panels show the mean images for the clusters to which the marked ion images are assigned. Similar to the lymph node dataset, the neural pipeline shows better cluster assignments than the regular pipeline, with mean cluster images that show a closer match to the assigned cluster images.

**Figure 7.**
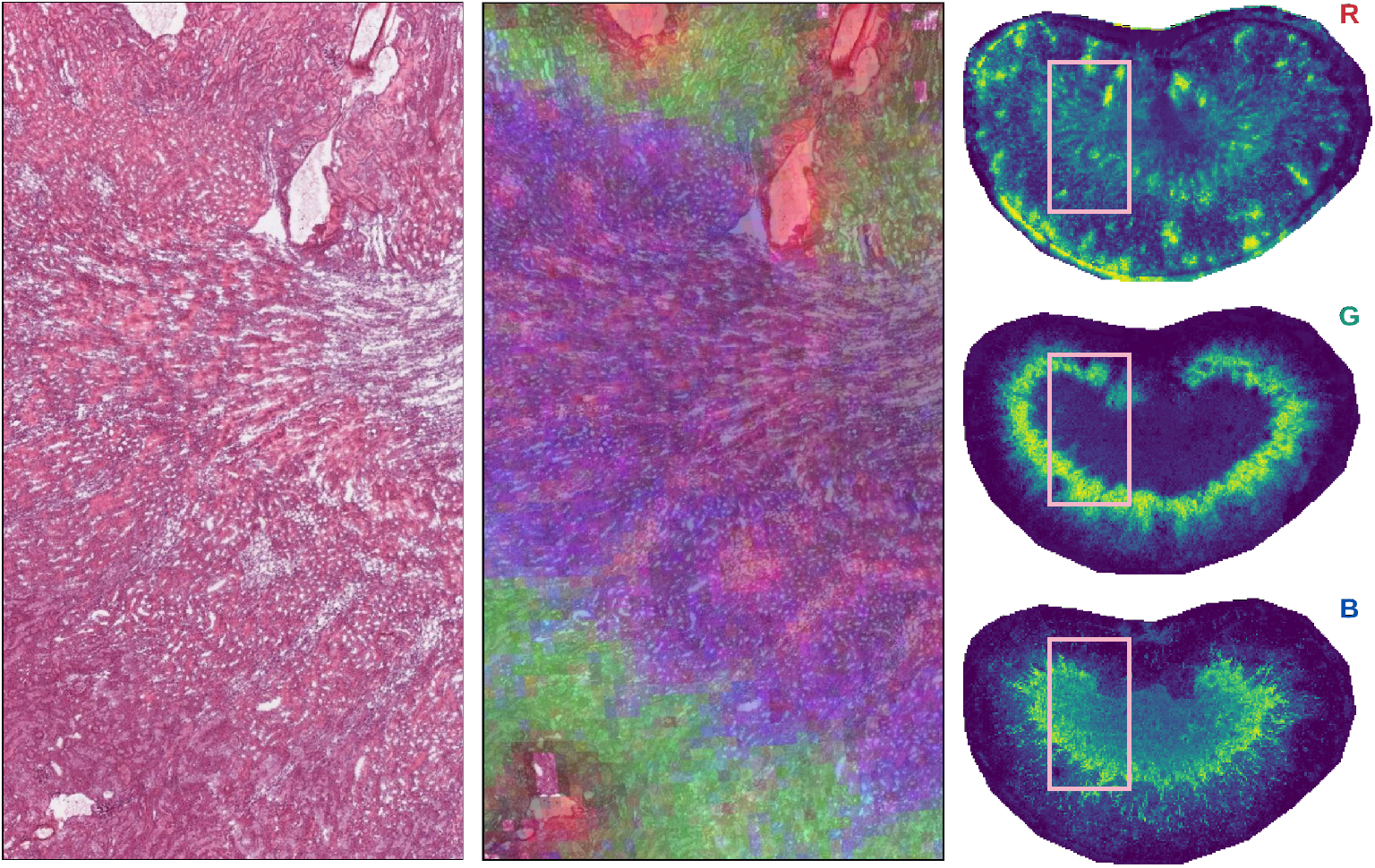
(Left) H&E stained microscopy image of neighboring kidney section. (Right) Three example mean cluster images for the neural U-D pipeline. (Center) Composite image combining the three mean cluster ions overlaid onto the microscopy. The co-localization of morphological patterns in the microscopic image and the spatial expressions in these mean cluster images corroborates the biological relevance of the clustering results.

### k-means clustering results

Finally, we apply k-means to the regular and neural ion images. When testing different numbers of clusters on the regular ion images, we run into the same issues as we did with clustering the regular ion images in the lymph node dataset when we increase the number of clusters, namely that we get a lot of clusters with a single ion image, due to the high sparsity of the data in the original feature space. For the best results in terms of detected meaningful clusters and RIR, we obtained around 20 clusters, similar to the regular U-D pipeline, which collects the most meaningful clusters and where there are no trivial clusters containing single ion images. Again, we do not encounter this issue with the neural pipeline, presumably due to a better packing of the data through neural feature extraction, and the lower dimensionality of the data (37, 365 pixels per regular ion image versus 2,048 dimensions in the neural ion images). As such we are able to explore more clusters via the neural pipeline. Similarly to the U-D results, we find clusters that better highlight localized structure in the neural pipeline, however, interestingly, the RIR for k-means is similar to that of U-D, showing that the relatively simple k-means performs surprisingly well on this dataset, compared to the much more involved U-D pipeline.

## Conclusions

We set out this work with the observation that the spatial information available in MSI is often under-utilized in its computational analysis, in part due to the fact that it is non-trivial to translate the complex spatial pattern recognition that humans perform on a daily basis into simple algorithms. The advent of deep learning, particularly convolutional neural networks and their derivatives, has significantly advanced the state of the art in computer vision, making it possible to capture high-level abstractions, learned through training complex models based on millions of images.

Moreover, a key strength of neural networks is the fact they can often be transferred between related tasks, such as object detection in computer vision. This enables the direct use of pre-trained neural networks to detect complex spatial patterns without having to train such a model from scratch for each application, which would often be intractable due to the required amount of data and computational resources. In this work we have used such a general-purpose pre-trained model as a general feature extractor to find spatial similarities between ion images in MSI data. Each ion image was fed through the network, and the resulting network interpretation, the neural ion image, was used for subsequent clustering of ion images.

We evaluated the clustering results of two different ion image clustering pipelines, namely the density-based DBSCAN clustering algorithm, combined with non-linear dimensionality reduction using UMAP, and the pervasive k-means clustering algorithm. In both pipelines, we compared regular ion images and neural ion images from two different MSI datasets, namely a human lymph node and a mouse kidney dataset.

All of the tested pipelines allowed for extraction of underlying spatial patterns in both datasets, showing insight in the underlying data. However, in all our experiments, the neural pipelines provided a better assignments of ion images, with more fine-grained clusters, and greater consistency in the spatial structures assigned to individual clusters, resulting in more representative mean cluster images. Specifically, our experiments indicated that using neural ion images enables the subsequent clustering pipelines to better account for localized, salient differences between ion images to obtain a fine clustering result, in contrast to the baseline methods which were less effective in identifying clusters with localized differences.

To quantify our results, we introduced a new metric called Relative Isotope Ratio, which measures the rate at which ion images of the same isotope are assigned to the same cluster, thus capturing biological relevance to a certain degree based on both spatial and spectral information. The newly introduced RIR metric quantitatively confirmed the merits of using neural ion images, thus corroborating our previously mentioned qualitative observations regarding improved clustering specificity.

In our experiments, neural networks greatly improved on traditional, global distance measures like cosine distance, whereas a neural network-based approach did not show significant improvement in performance over such distance measures in the work of Ovchinnikova et al. [29]. However, small differences in applied methodology can result in significant differences in outcome, and in our work, selecting a good physical patch size was instrumental in achieving good clustering results, something that was not explicitly mentioned by Ovchinnikova et al. It can be intuitively understood that an optimal patch size depends on the spatial patterns within the data itself, which, in the case of MSI ion images, has a clear link to the physical size of patterns within the underlying tissue, such as anatomical structures.

The proposed methodology can be readily used to incorporate an advanced form of spatial information into any MSI-focused machine learning pipeline, both supervised and unsupervised, and this without the need for large amounts of training data, or high computational needs. Furthermore, towards the future, a neural network could be trained specifically on MSI data, which we expect will further improve performance of our proposed pipeline.

## Acknowledgments

Wanqiu Zhang was supported by a Baekeland PhD grant from Flanders Innovation and Entrepreneurship (VLAIO). KU Leuven: Research Fund (projects C16/15/059, C3/19/053, C32/16/013, C24/18/022), Industrial Research Fund (Fellowship 13-0260) and several Leuven Research and Development bilateral industrial projects, Flemish Government Agencies: FWO (EOS Project no 30468160 (SeLMA), SBO project S005319N, Infrastructure project I013218N, TBM Project T001919N; PhD Grants (SB/1SA1319N, SB/1S93918, SB/151622)), This research received funding from the Flemish Government (AI Research Program). Bart De Moor and Wanqiu Zhang are affiliated to Leuven.AI - KU Leuven institute for AI, B-3000, Leuven, Belgium. VLAIO (City of Things (COT.2018.018), Innovation mandate (HBC.2019.2209), Industrial Projects (HBC.2018.0405)), European Commission: This project has received funding from the European Research Council (ERC) under the European Union’s Horizon 2020 research and innovation programme (grant agreement No 885682); (EU H2020-SC1-2016-2017 Grant Agreement No.727721: MIDAS)

